# The structure, redox chemistry and motor neuron toxicity of heterodimeric zinc-deficient SOD1-Implications for the toxic gain of function observed in ALS

**DOI:** 10.1101/2025.09.10.675162

**Authors:** Victor A. Streltsov, Katherine E. Ganio, Stewart D. Nuttall, J. Andres Hernandez, Cassandra N Dennys, Peter J. Crouch, Alvaro G. Estevez, Maria Clara Franco, Joseph S. Beckman, Blaine R. Roberts

## Abstract

A subset of familial cases of amyotrophic lateral sclerosis (fALS) are caused by mutations to copper, zinc superoxide dismutase (Cu, Zn SOD1). There are over 200 mutations to SOD1 that have been associated with fALS and the majority of these mutations are dominantly inherited. Thus, individuals are heterozygous and express both wild-type SOD1 and the mutant form of the protein. Paradoxically, when rodent models are produced that mimic the co-expression of wild-type SOD1 with mutant fALS SOD1 the motor neuron disease accelerates. Previously, we have shown that the loss of zinc from an SOD1 kills cultured motor neurons due to a gained, redox activity catalyzed by the active-site copper. Furthermore, motor neuron toxicity of zinc-deficient SOD1 is enhanced by wild-type Cu, Zn SOD1. Because SOD1 exists as a non-covalent dimer, the enhanced toxicity might result from stabilization of the heterodimeric interface between zinc-deficient SOD1 and Cu, Zn-SOD1. However, experimentation with the heterodimer is difficult because SOD1 subunits exchange in minutes. To better characterize the role of dimer stabilization on the enhanced toxicity of fALS mutant SOD1 by wild type SOD1, we genetically tethered a zinc-deficient SOD1 subunit with a Cu, Zn SOD1 subunit with a 16-residue linker. The x-ray structure of the tethered heterodimer shows that zinc-deficient subunit adopts a wild-type-like conformation and is not misfolded. The heterodimer intermediate also produced peroxynitrite from nitric oxide, and the tethered SOD1 was strikingly toxic to primary cultures of motor neurons. This work supports the concept that zinc-deficient SOD1 is a likely toxic intermediate in ALS. Furthermore, the wild-type allele in human familial-SOD1 ALS patients may physically contribute to the dominant inheritance of SOD1 mutations through heterodimer formation.

## INTRODUCTION

Amyotrophic lateral sclerosis (ALS) is an adult-onset disease involving the progressive death of lower motor neurons in the spinal cord and upper motor neurons in the brain stem and cortex, with consequent muscular paralysis (Rowland and Shneider, 2001). In 1993, thirteen missense mutations to the Cu, Zn superoxide dismutase (*SOD1)* gene were linked to the familial ALS (fALS) (Rosen et al., 1993). Structural analysis of the these variant showed an impact on the β-barrel fold and dimer interface (Deng et al., 1993). Since this discovery more than 200 dominant missense mutations and a dozen C-terminal frameshift and truncation mutations to the *SOD1* gene have been identified in ALS patients. Approximately 2-7% of ALS cases result from SOD1 mutations with the percentages varying in different regions around the world (Benatar et al., 2025). For many of the fALS SOD1 mutations, the structural changes produce subtle localized shifts in structure and many mutant SOD1s can retain full enzymatic activity. Virtually all fALS SOD1 patients have one remaining wild-type allele that produces enzymatically active Cu, Zn SOD1 protein.

Despite a lack of obvious structural or enzymatic effects, overexpression of fALS mutant SOD1 in mice and rats results in the progressive motor neuron death. These ALS models remain the most used animal model of human ALS. Overall, there is a consensus that *SOD1* mutations confer a toxic gain-of-function, rather than the loss of superoxide-scavenging activity. This is further supported by the acceleration of ALS when wild-type (WT) SOD1 is co-expressed with fALS SOD1 in transgenic animal models (Jaarsma et al., 2000; Witan et al., 2008). Accelerated death by WT SOD1 is also observed in motor neuron cultures isolated from fALS SOD1 mice (Garner et al., 2010).

We and others have hypothesized that the intermediates in the SOD unfolding process provide the foundation for toxicity that may explain the gain-of-function (Rakhit et al., 2004; Tiwari and Hayward, 2005). Indeed, careful biophysical studies have shown that the mutations to SOD1 result in a subtle decrease in protein stability affecting the dimer interface, and the degree of destabilization correlates with disease severity (Wang et al., 2008). Many fALS SOD variants are known to decrease in zinc binding affinity (Crow et al., 1997; Lyons et al., 1996; Smirnova et al., 2022; Souza et al., 2019), that is associated with a loss of dimer stability (Kumar et al., 2017). Loss of zinc has one of the most dramatic effects on the quaternary structure, distorting the dimer interface by 9° and disordering the electrostatic and zinc-binding loops (Roberts et al., 2007).

The absence of zinc results in disorder of the electrostatic loop, opening the narrow ∼4Å-wide active site channel that protects the reactive copper. The tight-fitting active site channel normally allows only superoxide to reach the mostly buried copper. One manifestation of the widening allows copper to become reduced to Cu(I) by low molecular weight reductants ∼3000 times faster than Cu, Zn SOD1 and thereby produce superoxide through the reduction of oxygen (Estevez et al., 1999). In the presence of a low concentration of nitric oxide, zinc-deficient SOD1 catalyzes the formation of peroxynitrite that activates a cell-death pathway leading to the death of primary motor neurons and makes astrocytes reactive (Barbeito et al., 2004; Estevez et al., 1999). The death pathway in motor neurons involves the nitration of HSP90 and activation of the ATP-dependent P2X7 receptor (Franco et al., 2013; Gandelman et al., 2013; Gandelman et al., 2010).

We have previously shown that the toxicity of zinc-deficient SOD1 in vitro is enhanced by the addition of WT Cu, Zn SOD1 (Garner et al., 2010). The increase in toxicity has been attributed to increased stabilization of zinc-deficient SOD1 through the dimer interface with Cu, Zn SOD1. Indeed, the disulfide bond of constitutively zinc-deficient SOD1 is less susceptible to reduction in vitro in the presence of WT Cu, Zn SOD1 and more resistant to aggregation (Garner et al., 2010; Roberts et al., 2007). We hypothesized that the Cu, Zn holo form of the enzyme structurally stabilizes the zinc-deficient SOD1 subunit and thereby protects this subunit from losing copper. To test this hypothesis, we made a tethered heterodimeric protein to prevent swapping of SOD1 subunits. The carboxyl-terminus of WT like Cu, Zn SOD1 (C111S) was tethered with a 16-residue linker comprised of glycine, serine, and alanine residues to the amino-terminus of zinc-deficient SOD1 (D83S/C111S) to produce a single chain protein modeling the heterodimer (Fig. 1). The linker was chosen from the well-known approach used for tethering heavy and light antibody chain fragments to create recombinant single-chain antibodies (Holliger et al., 1993). The x-ray structure, metal binding properties, redox activity, and toxicity to primary motor neurons were assessed using the tethered SOD1 heterodimer to determine how the dimer interface might contribute to toxicity.

**Figure 1.**
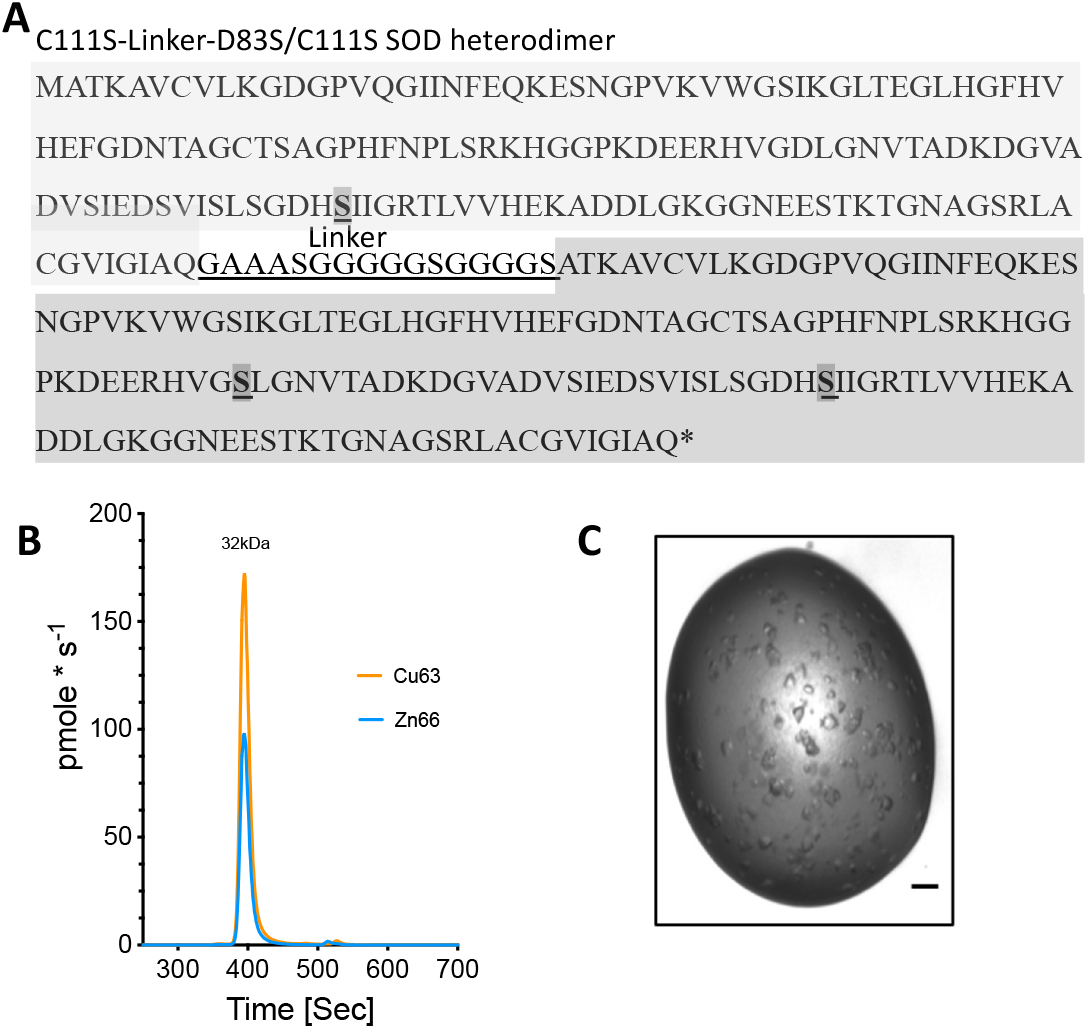
Tethered SOD1 sequence. Cloning, expression, and crystallization of C111S-D83S/C111S SOD1 heterodimer (Het-SOD1). (**A**) The SOD1 heterodimer is a single polypeptide chain of ∼32.7 kDa comprising tandem SOD1 protein monomers connected by a 16-residue flexible linker sequence (bold underlined). The darker shade with bold and underline indicate the location of mutations. The N-terminal monomer is wild type-like with a C111S mutation for better expression and solubility and the second constitutively zinc-deficient D83S/C111S C-terminal SOD1. (**B**) Size exclusion inductively coupled plasma mass spectrometry quantitation of Cu and Zn content. (**C**) Photograph of the crystals used for structural determination.

## MATERIALS AND METHODS

### Cloning and protein expression

The gene sequence for the wild-type-like SOD1 (C111S) tethered to zinc-deficient SOD1 (D83S/C111S) heterodimer (Het-SOD1) is shown in Fig 1. The D83S mutation alters one of the zinc ligands to produce a constitutively zinc-deficient SOD1 (Bahadorani et al., 2013; Estevez et al., 1999). Both subunits also contained the thermostable substitution at cysteine 111 to serine (C111S), which is known to be reactive in vitro and able to bind metals and complicate metal binding studies (Parge et al., 1992). The C111S mutation also improves expression of soluble SOD1 in *E. coli* (Garner et al., 2010; Leinweber et al., 2004). The sequence including 5’ *Nde*I and 3’ *Bam*H1 sites was synthesized at Geneart AG (Regensburg, Germany, www.geneart.com), sub-cloned into the expression vector pET3d (Novagen) and the sequence validated by complete sequencing. This wild-type-like SOD1 C111S + zinc-deficinet SOD1 D83S/C111S tethered heterodimer (Het-SOD1) was expressed in *E. coli* strain BL21(DE3) pLysS. Briefly, a single colony was cultured overnight in 300 mL Luria broth (LB) containing 0.1 mg/mL ampicillin at 37C and 180 rpm. The inoculate was added to three flasks, each containing 1 L of LB media and 0.1 mg/mL ampicillin and grown at 37 °C at 150 rpm to an OD_600_= 0.75. Protein expression was induced by addition of 1 mM (final) isopropyl-b-D-thiogalactoside. To obtain Cu, Zn-bound protein, CuSO_4_ (0.2 mM final) and ZnSO_4_ (0.1 mM) were added 1 h post-induction, the temperature decreased to 24°C, and expression continued for 16 h. Bacteria were pelleted by centrifugation (10,000 × g for 20 min) and stored at -20°C prior to processing.

### Protein purification of tethered WT SOD1+D83S

Bacteria were lysed by thawing with 1X tris-buffered saline (TBS, 50 mM Tris pH8.0, 150 mM NaCl) at a ratio of 0.5 mL of 1X TBS/g of pellet). DNase I bovine pancreas (3 mg) was added followed by incubation at 37 °C for 2 h. Lysates were clarified by centrifugation (10,000 g for 20 min) at room temperature (RT), the supernatant adjusted to 35% ammonium sulfate followed by a further clarification by centrifugation as above. The solution was then brought to a final concentration of 70% ammonium sulfate followed by centrifugation as above. The supernatant was applied to a Phenyl Sepharose column (GE Healthcare, 5×50 mm GL) equilibrated in 2 M NH_4_SO_4_ + 150 mM NaCl and a linear gradient over 10 column volumes from 0 – 100 % buffer B (50 mM KP_i_, pH 8). SOD1 positive fractions, determined by SDS-PAGE, were pooled and exhaustively dialyzed against 10 mM KP_i_, pH 8 at 4°C. Dialyzed material was loaded on a 5/50 GL Mono Q anion exchange column (GE Healthcare, 5×50 mm) with an elution gradient from 10 mM KP_i_, pH 8 to 10 mM KP_i_+ 100 mM NaCl, pH 8 over 20 column volumes. Fractions from the anion exchange column containing SOD1 were pooled and concentrated using 5K MWCO Millipore Amicon Ultra-50 Centrifugal Filter Units. The concentrated material was loaded on to a 10/300 GL Superdex 75 size exclusion column developed with 1X phosphate buffered saline (PBS). SOD1 fractions corresponding to the heterodimer molecular mass of 32kDa were pooled and concentrated using 3K MWCO Millipore Amicon Ultra-50 Centrifugal Filter Units. We also observed a small fraction of a tetrameric SOD species (64kDa) that we ascribe to formation of a dimer of two tethered heterodimers in the crude lysates that was purified away (data not presented). Protein concentration was calculated using an extinction coefficient of 5810 M^-1^ cm^-1^ at 280 nm, which is based on the absorbance calculated for the denatured SOD protein sequence (Gill and von Hippel, 1989).

### Peroxynitrite generation assay

The boronate-based fluorogenic probe, coumarin-7-boronic acid (CBA), was used to detect peroxynitrite formed during the production of superoxide via SOD1 re-oxidation in the presence of nitric oxide. CBA was a kind gift from Drs. Zielonka and Kalyanaraman at the Medical College of Wisconsin. CBA was dissolved in dimethylformamide and stored at -80 °C until use. Re-oxidation experiments were performed at 37 °C in 20 mM HEPES buffer, pH 7.4 containing 100 µM CBA, 5 µM ascorbate, 10 µM SOD, and 5 µM PAPA-NONOate as steady-state source of nitric oxide. Catalase (100 U/mL) was added to prevent CBA oxidation due to H_2_O_2_. Fluorescence of coumarin was measured at excitation/emission wavelengths of 330/455 nm using a Gemini fluorescence microplate reader (Molecular Devices, Sunnyvale CA). Standard curves for quantifying peroxynitrite generation were determined by reacting a known amount of peroxynitrite with CBA. Peroxynitrite was synthesized from nitrite and hydrogen peroxide using previously described methods (Robinson and Beckman, 2005). The concentration of peroxynitrite was determined by measuring absorbance at 302 nm (ε= 1,700 M^−1^ cm^−1^). Peroxynitrite working stocks 100x were prepared in cold, degassed water and a small aliquot was immediately added to degassed 20 mM HEPES buffer, pH 7.4 containing 100 µM CBA and 100 U**/**mL catalase. Relative fluorescence was converted to peroxynitrite concentration using non-linear regression fitted as a single exponential. Standard curves for CBA were applied to experiments to quantify peroxynitrite generation.

### Size-Exclusion Chromatography with Inductively Coupled Plasma Mass Spectroscopy (SEC-ICP-MS) metal assay

The metal status of the SOD1 heterodimer was determined using a SEC column (BioSEC3 4.6 × 300mm, Agilent) connected directly to an ICP-MS (Agilent 7700) as previously described(Lothian and Roberts, 2016). Breifly, the column flow rate was 0.4 mL*min^-1^ with 200 mM ammonium nitrate pH 7.7-7.8. The column temperature was maintained at 30°C using a thermostatically controlled column compartment. The ICP-MS was run under standard multi-elemental conditions and bovine SOD1 (Sigma) was used as a molecular mass standard and a standard for copper and zinc. A standard curve for copper and zinc was generated using bovine SOD1 (200 ppb Cu and zinc) injection volumes ranged from 3-30 µL. The integrated area for SOD1 was determined for copper and zinc traces and the response for the ICP-MS in picograms/sec was determined by dividing picograms of copper or zinc injected on the column over the integrated peak area in counts per second (CPS). The picograms/count constant was then used to convert CPS to picograms of metal per second.

### ESI-MS determination of mass and validation of metal status

The metal content and mass of the D83S+WT heterodimer SOD1 was determined by native electrospray mass spectrometry using a size exclusion column (AdvanceBio SEC 200Å, 1.9µm 2.1 × 50mm) coupled to a QTOF with Jet stream source (Agilent 6545XT). The column was developed with 100 mM ammonium acetate pH 7.0. Instrument setting was as described in Supplemental Table 1. Data were collected in profile mode and deconvoluted with Bioconfirm software (Agilent Technologies) with the mass range 600-5000, mass step of 0.05 and baseline factor 7.00. The observed deconvoluted mass for the heterodimer with metals bound was 32,759.14 g/mol (5.5 ppm mass error, most probable mass 32,758.96). The most probable mass for the heterodimer C111S+D83s/C111S SOD1 was calculated with iMass application (v1.4, Mobile Science Apps.).

### Heterodimer SOD1 crystallization

Crystallization screening was performed at the CSIRO Collaborative Crystallization Centre (www.csiro.au/c3). C111S+D83S/C111S SOD1(Het-SOD1) concentrated to ∼10 mg/mL in PBS buffer was set up as 0.4 μL sitting drops at 20°C against a customized 96-well screen, based upon successful previous crystallization conditions for SOD1 (Roberts et al., 2007), heavily biased toward 2-3 M ammonium sulfate or sodium malonate at pH range 4.5-7. Crystals grew under a variety of conditions; the best diffracting crystals were 100-200 µm in size grown in sodium malonate for 7 – 21 days.

### X-ray crystallography

X-ray diffraction data sets for single crystals of Het-SOD1 in well solution plus *ca* 15% (v/v) glycerol as cryo-protectant were collected at 100 K using the MX2 beamline at the Australian Synchrotron, a micro-focused in-vacuum undulator beamline. The data sets were processed with HKL2000 (Otwinowski and Minor, 1997). Further data collection and processing statistics are given in **Table 1**. The locations of SOD1 monomers were identified in the asymmetric units by PHASER (McCoy et al., 2007) molecular replacement using the structure of SOD (PDB ID: 2R27) (Roberts et al., 2007) without including water molecules. Five independent heterodimers were identified in the asymmetric unit of the Het-SOD1 crystal. The structures with protein molecules alone were initially refined and visible parts of linker sequences plus water molecules were subsequently added. Occupancies of metal sites were carefully refined with constrained atomic B-factors. Iterative refinement and model building were conducted using REFMAC (Murshudov et al., 1997) and XFIT/MIFit (McRee, 1999). Progress of the refinement was monitored using the *R*_free_ statistics based on a test set encompassing 5% of the observed diffraction amplitudes (Brunger, 1992). Deposition codes and further structure refinement details are given in **Table 1**. The figure was produced using PyMol (Schrödinger Software).

**Table 1.** Data collection, processing and refinement. Values for the outer shell are given in parentheses.

| D83S-C111S SOD heterodimer PDB ID 6DTK |  |  |
| --- | --- | --- |
| *Data collection* |  |  |
| Diffraction source | Australian Synchrotron MX2 |  |
| Wavelength (Å) | 0.95370 |  |
| Temperature (K) | 100 |  |
| Detector | ADSC Quantum 315r |  |
| Space group | $C222_1$ | |
| $a, b, c$ (Å) | 162.70, 201.67, 143.83 | |
| $\alpha, \beta, \gamma$ (°) | 90, 90, 90 | |
| Mosaicity (°) | 0.3 |  |
| Resolution range (Å) | 47.57–2.00 (2.05–2.00) |  |
| No. of unique reflections | 158995 (15780) |  |
| Completeness (%) | 99.9 (98.9) |  |
| Redundancy | 14.2 (13.4) |  |
| $\langle I/\sigma(I) \rangle$ | 25.25 (2.47) | |
| $CC_{1/2}$ | 0.6 | |
| $R_{\text{r.i.m}}$ | 0.086 (0.871) | |
| Solvent content (%) | 66.27 |  |
| Matthews coefficient (Å<sup>3</sup>/Da) | 3.65 |  |
| Overall $B$ factor (Å<sup>2</sup>) | 38.62 | |
| *Refinement* |  |  |
| No. of reflections, working set | 150990 (2590) |  |
| No. of reflections, test set | 10937 (576) |  |
| Final $R_{\text{cryst}}$ | 0.156 (0.225) | |
| Final $R_{\text{free}}$ | 0.190 (0.243) | |
| No. of non-H atoms |  |  |
| Protein | 11297 |  |
| Ion | 29 |  |
| Solvent | 7 |  |
| Water | 1357 |  |
| Total | 12690 |  |
| R.m.s. deviations |  |  |
| Bonds (Å) | 0.13 |  |
| Angles (°) | 1.36 |  |
| Average $B$ factors (Å<sup>2</sup>) | | |
| Protein | 38.8 |  |
| Ion | 34.8 |  |
| Solvent | 39.5 |  |
| Water | 51.8 |  |
| Ramachandran plot |  |  |
| Most favoured (%) | 98.1 |  |
| Allowed (%) | 1.7 |  |

### X-ray Absorption Spectroscopy (XAS)

XAS spectra of frozen solution samples of <0.5mM concentration of wtSOD1, Het-SOD1 without and with 20-fold concentration of sodium ascorbate as a reductant were collected at the Australian synchrotron XAS beamline (1.9 T Wiggler). The beamline was equipped with liquid nitrogen (LN_2_) cooled Si double crystal monochromator (ΔE/E 1.5×10^-4^) with a Rh-coated focusing mirror to produce a focused X-ray beam with a harmonic content better than 1 part in 105. The incident and transmitted x-ray intensity was monitored using ionization chambers with a continuous stream of He gas. Fluorescence measurements were obtained using a 100-element LN_2_-cooled Ge detector (Canberra). Energy calibration was achieved by the simultaneous accumulation of a Cu foil spectrum (transmittance) where the inflection point of the first absorption feature was set to an energy of 8980.4 eV. Ice formation was inhibited by addition of glycerol (∼15%) to samples immediately prior to their injection into the 40 µL cavity of polycarbonate cells (2 mm × 2 mm × 10 mm) with Kapton (Goodfellow Cambridge, Cambridge, UK) front and back windows. Samples were frozen and stored in liquid N^2^ until transfer to the beamline closed-cycle pulse tube He cryostat (“Optisat”, Oxford Instruments).

A series of Zn (9659 eV) and Cu (8979 eV) K-edge XANES (X-ray Absorption Near Edge Spectroscopy) measurements scans up to k = 10 Å^-1^ were obtained from samples in a fluorescence mode at 5-10K. Radiation damage of samples was tested by quick XANES measurements from the same sample position with 30 min exposure intervals. The spectra recorded from each sample position were averaged to obtain the final spectra. The XANES spectra were pre-processed by the software package SAKURA at the Australian synchrotron XAS beamline. Data points affected by monochromator glitches were also removed. Edge step normalization for each spectrum was performed by subtracting the pre-edge and post-edge backgrounds in program ATHENA (Ravel and Newville, 2005) based on the IFEFFIT library on numerical and XAS algorithms (Newville, 2001).

### Motor neuron culture and survival assays

Motor neurons were purified from E15 rat embryos by a combination of density centrifugation and immunoaffinity using an antibody to the p75 low affinity neurotrophin receptor as previously described (Raoul et al., 2002). Purified motor neurons were plated in 96 well plates at a density of 500 cells per well in Neurobasal media supplemented with B27, b-mercaptoethanol, glutamine, glutamate, horse serum, brain-derived neurotrophin factors (BDNF), glial-derived neurotrophic factor (GDNF) and cardiotrophin-1. After 24 h in culture, SOD1 was delivered utilizing the membrane permeant carrier agent Chariot (Active Motif, Carlsbad, CA). SOD1 was diluted in H^2^O at a final concentration of 5 µg/mL at room temperature. SOD1 was incubated at room temperature before mixing with Chariot and incubated for additional 30 min for complexes formation. The medium was aspirated and replaced by Opti-MEM transfection medium (Invitrogen) containing the mixture Chariot and SOD1. The cells were incubated for 1 h, at which time 100 µL of Neurobasal medium containing twice the concentration of supplements and penicillin-streptomycin plus or minus trophic factors was added to each well and further incubated for 24h. Motor neuron survival was determined by high throughput image capture and analysis in 96-well plates using a Runner™(Trophos, Marseilles, France) as reported previously (Franco et al., 2013; Garner et al., 2010). Survival was standardized between experiments by considering the survival in the presence of trophic factors as 100%.

## RESULTS

### Protein expression, characterization, and crystallization

The heterodimer SOD1 (Het-SOD1) composed of wild-type like SOD1 (C111S) tethered to zinc-deficient SOD1 (D83S/C111S) expressed well in *E. coli* at yields greater than 5 mg/L (Fig. 1A). Following purification, size exclusion, inductively coupled mass spectrometry (SEC-ICP-MS) was used to determine metal content (Fig. 1B). The observed ratio of 1.76 Cu per Zn in the SOD1 heterodimer is slightly less than the expected ratio of two coppers and one zinc per mole of heterodimer SOD1.

Native mass spectrometry confirmed that the major Het-SOD1 protein had two coppers and one zinc (Fig. 2B). This peak had an average mass of 32,759.0 Da, while the theoretical average molecular weight is 32,760.0 Da for the heterodimer (C_1395_H_2220_N_422_O_467_S_6_Cu_2_Zn_1_), yielding a mass error of -30.5 ppm, which is excellent agreement. A less intense peak with a mass of 32,695.2 Da differed from the main peak by a mass of 62.8 Da, which is consistent with the loss of one additional metal. Other small peaks are consistent with one or two sodium adducts as well as a minor peak corresponding to the loss of two metals. Together, the ICP-MS and native results show that the Het-SOD1 protein was predominantly one peptide chain containing two coppers and one zinc, with a smaller fraction most likely missing a copper based on the Cu/Zn ratio determined by ICP-MS.

### Motor neuron survival

Previously, we have shown that SOD1 protein can be delivered intracellularly with *Chariot™* (active Motif, Carlsbad, CA)(Sahawneh et al., 2010). Consistent with prior results, zinc-deficient D83S SOD1 in the presence of trophic factors activated cell death by a nitric oxide-dependent oxidative mechanism that resulted in 53±7% survival of motor neurons in 24 hours (p-value<0.001). In contrast, Cu, Zn bound-SOD1 did not diminish or increase survival (Fig. 3). However, addition of an equal concentration of Cu, Zn bound SOD1 with zinc-deficient D83S SOD1 decreases survival to 19±6.1%. The Het-SOD1 heterodimer further decreased motor neuron survival to 10±6.3%.

**Figure 3.**
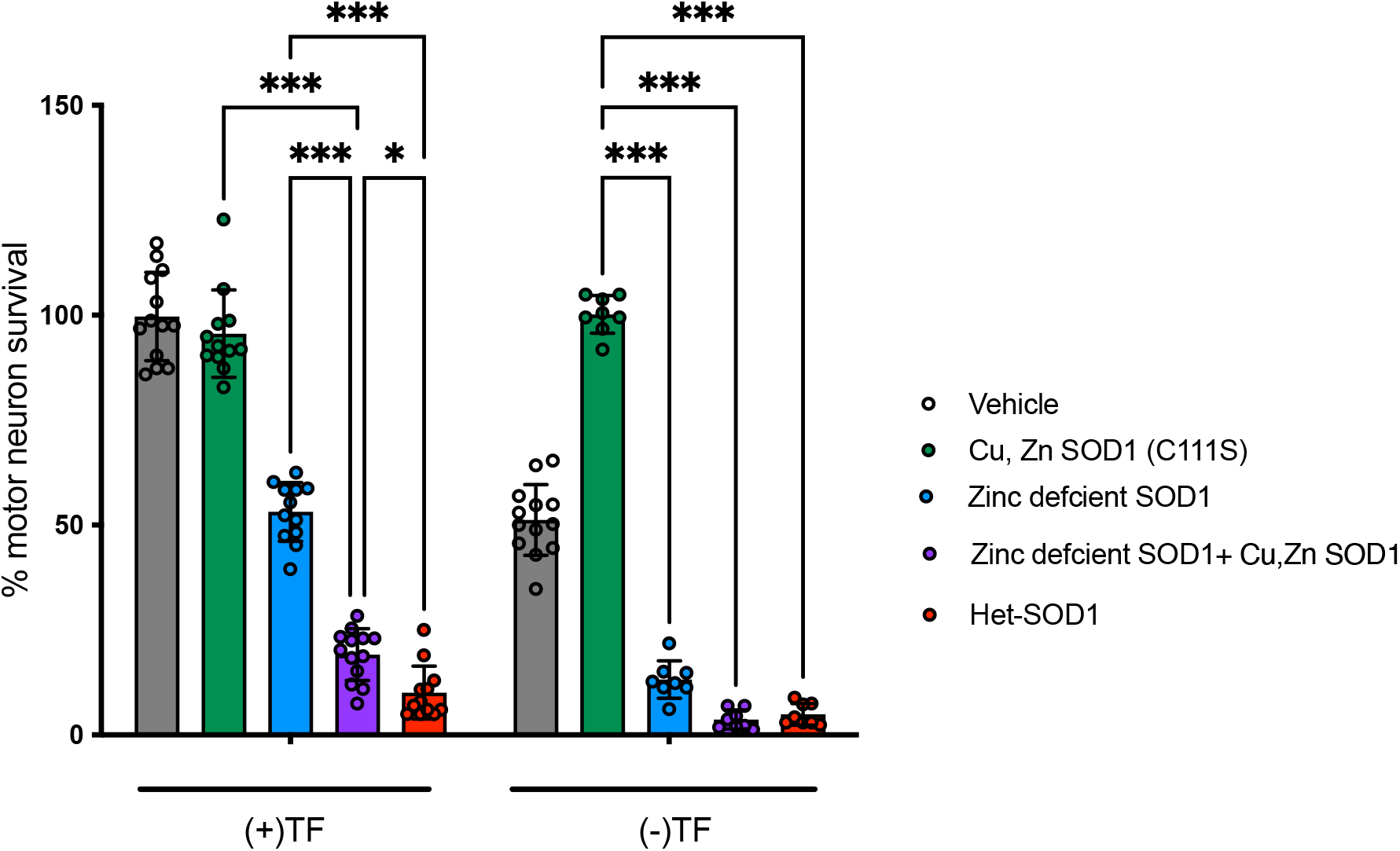
Motor neuron survival assays. Motor neuron survival was determined after administration of vehicle only (black bar vehicle, n=13 each condition), C111S-SOD1 (grey bar WT, n=12 TF, n=8 TFD), zinc deficient D83S/C111S SOD1 (green, n=12 TF, n=8 TFD), a 50/50 mixture of zinc-deficient D83S/C111S-SOD1 and holo SOD1(C111S) (grey diagonal lines, Cu, Zn SOD1, n=13 TF, n=8 TFD) and the Het-SOD1 with the glycine serine linker (red bar D83S/C111S-C111S SOD1 heterodimer, n=12 TF, n=8 TFD). Cultures treated with the trophic factors BDNF, GDNF and cardiotrophin-1 (TF) and separately after trophic factor deprivation (TFD). Values are mean with standard deviation. Statistical analysis with two-way ANOVA with Tukey’s multiple comparison post-hoc test (*p-value 0.02, ***p-value <0.001).

Motor neurons intentionally deprived of trophic factors undergo cell death with 51±8.4% survival at 24 h. Consistent with prior results (Garner et al., 2010), delivery of Cu, Zn bound SOD1 strongly protected motor neurons (100.±4.6% survival). Zinc-deficient SOD1 (D83S) by itself further decreased survival to 13±2.4% and the co-delivery with Cu, Zn bound SOD1 further decreased survival to 3.5±2.3%. Tethered Heterodimeric SOD1 (zinc-deficient-holo, D83S/C111S+C111S SOD1) also resulted in a significant decrease in survival 4.9±2.6%. Thus, holo Cu, Zn SOD1 by itself protected motor neurons from trophic factor deprivation, but zinc-deficient SOD1 alone or as a tethered heterodimer decrease survival consistent with previous reports (Estevez et al., 1999; Garner et al., 2010).

### Peroxynitrite generation by zinc-deficient SOD

Because we have previously shown that zinc-deficient SOD1 activates motor neuron death by an oxidative mechanism requiring both nitric oxide and superoxide, we next compared the peroxynitrite generation activity of the tethered heterodimer (D83S-WT SOD1) to that of zinc-deficient SOD1. The coumarin boronate probe has proved to be more specific and sensitive assay for peroxynitrite generation than other fluorescent oxidation probes (Zielonka et al., 2012). Under the current assay conditions (Fig. 4), the probe slowly hydrolyzed at a rate equivalent to 8.3±0.2 nM.min^-1^ and this rate was only slightly decreased with the addition of 10 µM WT-Cu,Zn SOD1 to 6.8±0.1 nM.min^-1^. Zinc-deficient WT SOD1 produced peroxynitrite at an apparent rate of 25.6±0.3 nM.min^-1^ per µmol SOD1, while the D83S+WT heterodimer produced peroxynitrite at a slightly slower but similar rate of 23.5±0.3 nM.min^-1^.

**Figure 4.**
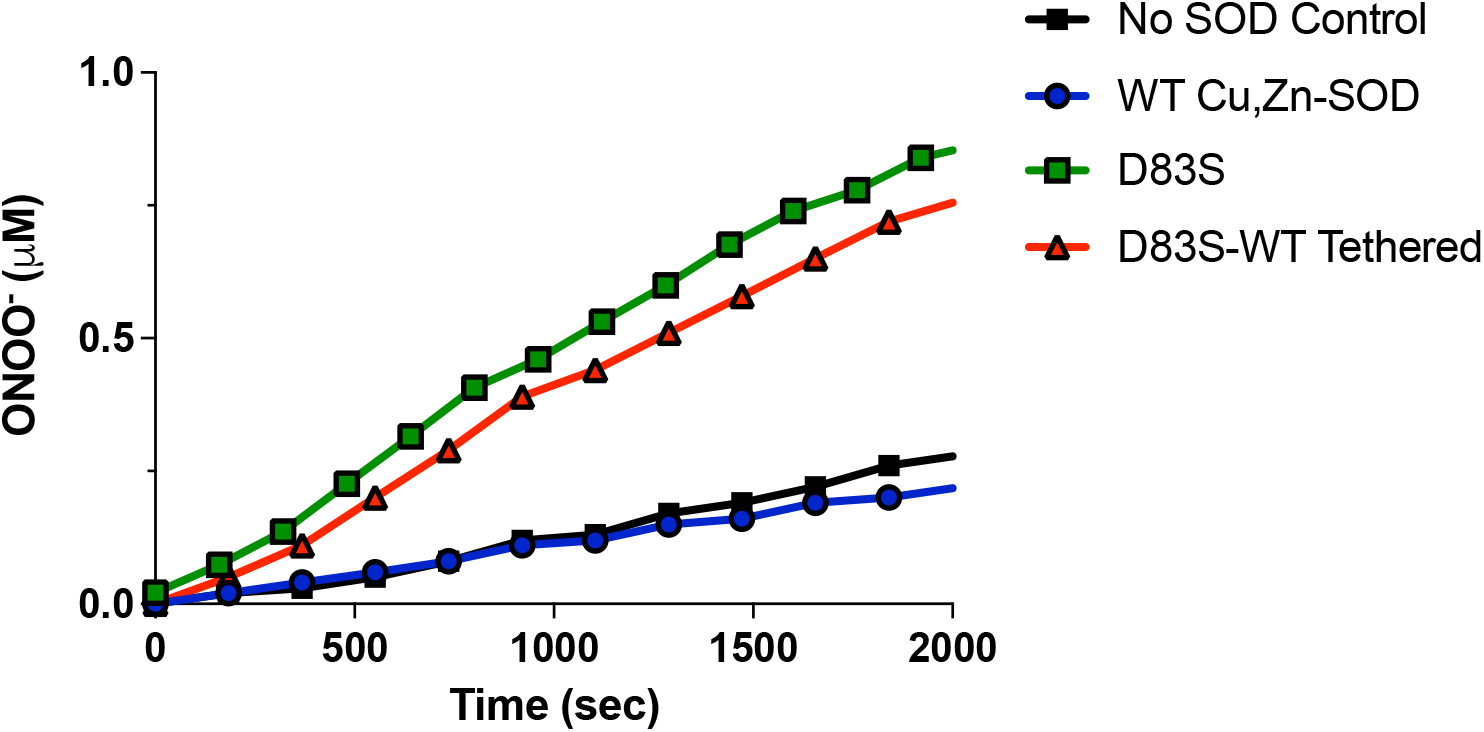
Peroxynitrite production due to re-oxidation of SOD. Preparations of Zinc-deficient WT SOD including wild type Cu, E SOD (green squares) and the D83S D83S+WT heterodimer (red triangles) are readily reduced via ascorbate to produced superoxide. In the presence of NO, superoxide reacted to form peroxynitrite, which was detected via oxidation of coumarin boronate assay. WT Cu,Zn SOD (blue circles) did not oxidize coumarin boronate any faster than the spontaneous hydrolysis compared to buffer control (black squares).

### X-ray crystal structure of D83S+WT heterodimer

Initial crystallization conditions for SOD1 homodimers had a marked preference for solutions rich in ammonium sulfate. Accordingly, a customized 96-well screen was designed around ammonium sulfate and sodium malonate. Multiple wells containing cuboidal crystals typically of 100-200 nm in length were found between 7 to 21 days (Fig. 5).

**Figure 5.**
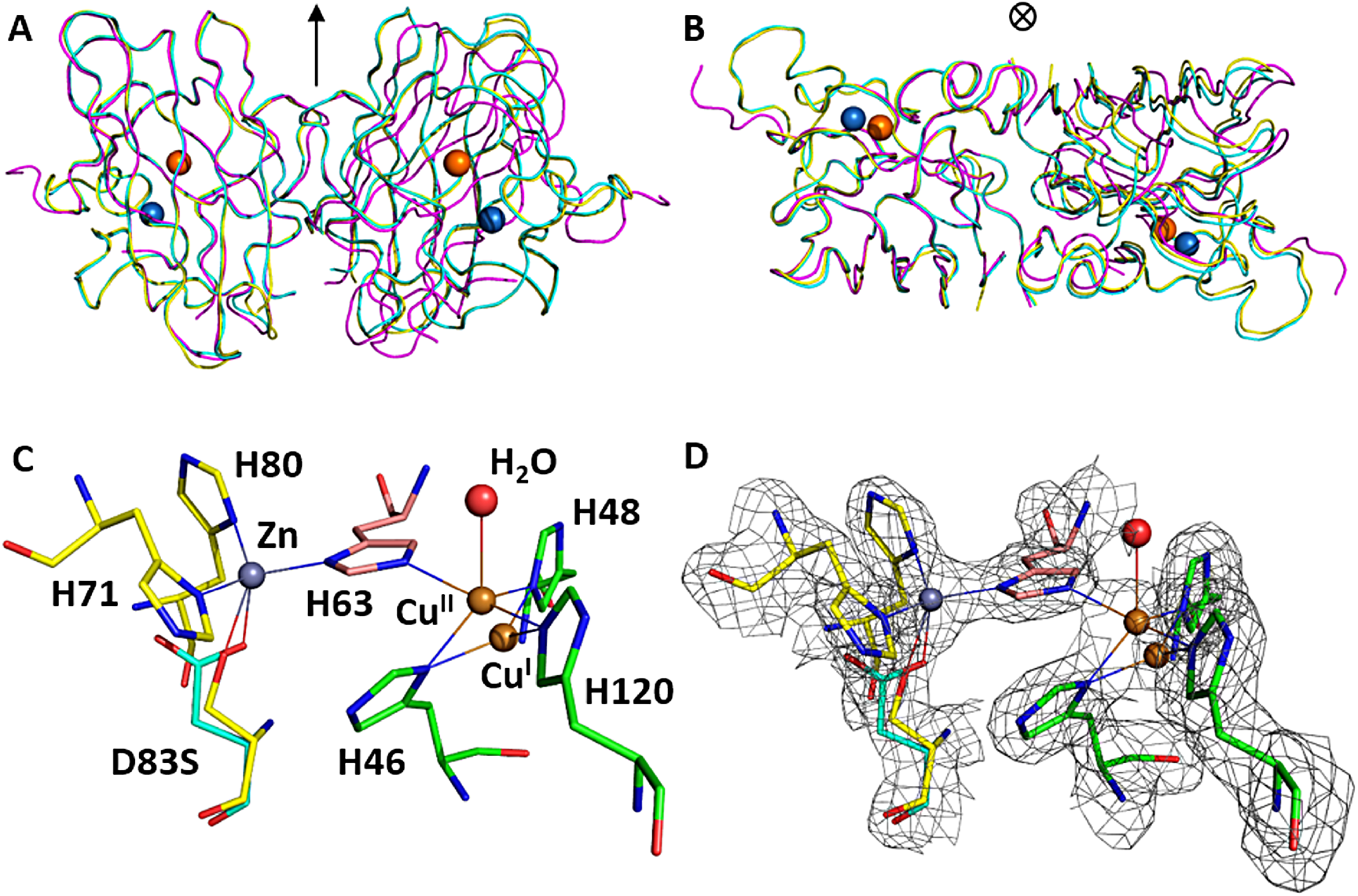
Structure of heterodimer D83S-C111S SOD1 (yellow, 6DTK), wild type SOD1 (1PU0) (cyan) and D83S mutant (2R27) (purple) are overlaid using one monomeric subunit A (left) to show perfect superposition of the second subunit B heterodimer and wild type and significant twist of the mutant monomer. **A)**. Front view and **B)** Top view. The approximate two-fold axis is shown by the arrow, Zn is in blue and Cu^II^ is in orange. **C)** Representative coordination environment of Zn and Cu in heterodimer D83S/C111S-C111S SOD1, and **D)** overlaid with the electron density map shown as a grey mesh contoured at 1.5σ within 1.6 Å of ligands.

The D83S+WT heterodimer SOD1 readily crystallized in a form (space group *C* 2 2 1 with a=162.7 Å, b=201.7 Å, c=143.8 Å). The asymmetric unit contained five SOD1 heterodimers with a high solvent content (66.3%), and is highly similar to the asymmetric subunits of WT SOD1 that have been previously reported (DiDonato et al., 2003; Parge et al., 1992). The structure was solved by molecular replacement which identified ten subunits, or five heterodimers. The 16-residue linker between monomers were partially visible in heterodimers and were built into available density. The structure was refined to 2.0 Å resolution with final R/R^free^ factors of 0.156/0.190 (Table 1) with well-ordered parts and coordinate accuracy of ∼0.1 Å. The ten monomeric subunits were refined individually without the use of non-crystallographic symmetry restraints to assess subtle structural changes between the heterodimer and wild-type dimer. Overall, these results suggest insignificant structural changes compared to the wild type variants and the linker region had minimal impact on the overall SOD1 structure.

The basic fold of zinc-deficient D83S subunit remained unchanged. The main chain conformations for the 153 residues were well defined as previously characterized (Fig. 5). The average RMSD between all five heterodimers in the asymmetric unit was 0.41 Å. The average RMSD between the D83S-C111S heterodimer (6DTK) and the homodimer of the wild type SOD1 (PDB ID: 1PU0) (DiDonato et al., 2003) was 0.30 Å. Comparison of the Het-SOD1 with the homodimeric zinc-deficient D83S SOD1 structure (PDB ID: 2R27) (Roberts et al., 2007) produced a RMSD of 0.66 Å. Similarly, the RMSD of alignment between wild type SOD1 (1PU0) and zinc-deficient SOD1 (2R27) was 0.67 Å. Hence, the structure of D83S+WT SOD1 heterodimer was more similar to wild-type Cu,Zn homodimeric structure than the Zn-deficient homodimeric structure. This is most pronounced in the strcutre of the electrostatic and zinc-binding loops were are ordered in the heterodimer structure (Fig. 5).

Furthermore, the C-alpha backbones of heterodimeric subunits exhibit near two-fold symmetry with the average RSMD of 0.25 Å, further supporting the closer similarity to the wild-type SOD. Previously, we showed that the loss of zinc results in local unfolding of the electrostatic and zinc binding loops. This was associated with a shift in the quaternary structure that is expressed as a 9.9° rotation of the dimer due to an opening of the dimer interface (Roberts et al., 2007). Due to the inherent link of the dimer interface with the zinc-binding loop, we proposed that a heterodimer consisting of one monomer of Cu, Zn bound-SOD1 with one monomer of zinc-deficient SOD1 was expected to correct the shift in the quaternary structure. This was based on the observations that mixing zinc-deficient SOD1 with Cu, Zn bound-SOD1 resulted in protection of the disulfide bond to reduction and resistance to protein aggregation (Garner et al., 2010). The crystal structure presented here demonstrates that the heterodimer corrects the quaternary structure of the dimer with a rotation angle between the two subunits of only -0.3° different compared to wild type enzyme (1PU0) (Figure 3A, B). This angle was nearly identical to the wild type SOD1 (1PU0) (DiDonato et al., 2003; Tainer et al., 1983).

### Active site structure

The electron density map confirmed the presence of mutated side chain D83S and C111S. However, the electron density of the D83S side chain in the zinc coordination spheres showed a mixed population of the mutant sites (Fig. 5 C, D). This might possibly be due to crystal averaging of subunits, because the heterodimers exhibit near two-fold symmetry. Therefore, the mutant site was specifically refined using mixed fractional population of two residues D83 and S83 while keeping the atomic B-factor of two amino acids restrained to similar values. The population of zinc at this site was also refined with a fixed, physically reasonable atomic B-factor. The resulting deficiency of zinc site in the range of 0.3-0.8 (0.64 on average) was proportional to the population of Serine-83 in the mutated site. The quasi-tetrahedral coordination of zinc with D83 ligand remained similar to that found in WT SOD1 with the distance between zinc and D83 of 1.97-1.99 Å and with histidines H63, H71 and H80 at 2.0-2.1 Å. The zinc coordination site varied in subunits with mutant D83S ligand being at 2.3-3.5 Å and with occasional additional waters located 2.3 Å from zinc.

The electron density of the copper site showed that fractions of copper can be modeled at alternate positions shifted by 1.1 Å in average over the structure (Fig. 3 C, D). The coordination of copper in oxidized (Cu^II^) site involved residues: His46, His48, His120 and His63 that is the bridging ligand for both copper and zinc. An additional one or two waters at the average distance of 2.5 Å resulting in five- or even six-coordinated site. The bridging His63 – Cu^II^ distance tends to be somewhat longer of ∼2.2 Å and the increased chemical-reactivity measured in the peroxynitrite generation assay stance tends to be somewhat longer of ∼2.2 Å and the imidazole plane is slightly tilted relative to the Cu-N bond. The coordination of copper in reduced (Cu^I^) site was approximately trigonal with binding residues: His46, His48, His120. Importantly, the population of Cu^I^ site always refined to be significantly greater than that of the Cu^II^ site with the ratio of Cu^I^ / Cu^II^ of 2.61 averaged over all heterodimers in the unit cell. The ratio of total copper to zinc was of 1.57, which agrees with the SEC-ICP-MS estimate of ∼1.78. Even though all of the C57-C146 disulfide bonds were in place, indicating absence of marked radiation damage, the proportion of Cu^I^ vs Cu^II^ could have been influenced by radiation-induced reduction of copper by a high flux of X-ray synchrotron radiation during data collection (Burmeister, 2000; Deng et al., 1993; Hart et al., 1999; Stroppolo et al., 1998; Weik et al., 2000). This was confirmed by X-ray absorption spectroscopy as described below. The bound copper redox potentials in SOD1 are within the range of x-ray absorption energy changes that occur due to alterations in protein conformations (Hart et al., 1999; Hart et al., 1998).

In contrast to the highly disordered zinc-binding section of loop IV (residues 68-78) and adjacent residues 132-139 of the electrostatic loop VII reported in the zinc-deficient homodimer structure (Roberts et al., 2007), the residues in these loops assumed their wild type conformation, but with higher atomic displacement parameters (B-factors) compared to the rest of the structure in D83S subunit. Therefore, the active-site channel was less disrupted compared to Zn-deficient homodimer (Roberts et al., 2007). The conformations of conserved Arg143 in heterodimer remained similar to those seen in wild-type Cu, Zn SOD1 and the disulfide subloop was not significantly distorted and shifted.

### X-ray Absorption Near-Edge Spectroscopy (XANES)

To investigate the redox status and coordination of the Cu and Zn metals and whether photoreduction played a role in our crystallography experiments at the synchrotron source, the X-ray absorption edges of frozen solution samples were collected at 10K. Previous reports demonstrated that metal coordination geometry is unaffected by the solution or by the crystalline state of SOD1 (Ascone et al., 1997). XANES data were obtained for Cu and Zn K edge for four samples: WT SOD1, WT SOD1 reduced with ascorbate, D83S-C111S heterodimer with and without reduction by ascorbate (Fig. 6). The experiments were performed at the Australian Synchrotron XAS beamline, where the photon flux was two orders of magnitude lower than in the MX2 beamline used for crystallography. No significant radiation damage or photoreduction was detected during irradiation, as judged by comparing individual XANES scans at the start and finish to the measurements.

**Figure 6.**
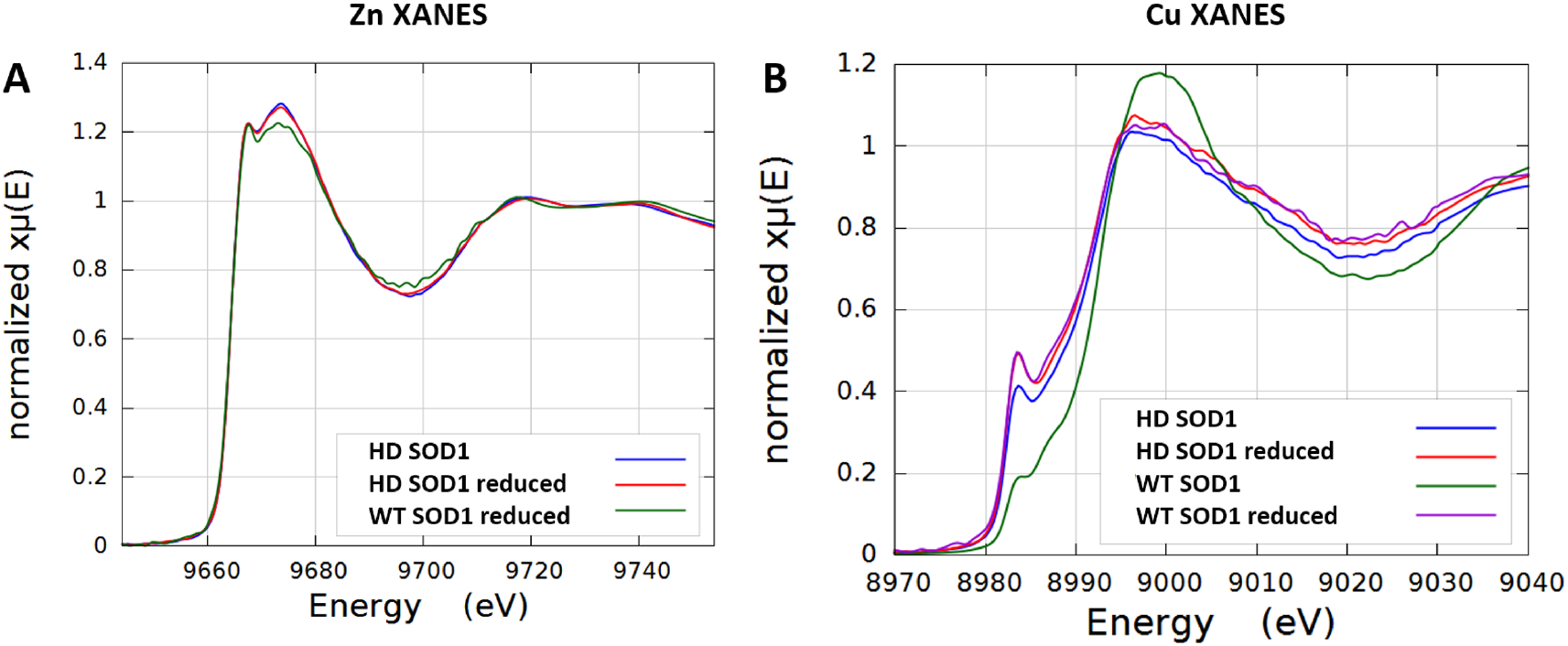
XANES spectra for (A) Zn K-edges of D83S-C111S SOD1 heterodimer (HD) (blue), reduced Het-SOD1 (red), and WT SOD1 reduced (green), and for (B) Cu K-edges of D83S+WT SOD1 heterodimer (HD) (blue), reduced HD SOD1 (red), WT SOD1 (green), and reduced WT SOD1 (purple).

The XANES spectra for Zn K-edges for D83S-C111S heterodimer in both its untreated and chemically reduced states (Fig. 6A) overlay well, supporting the structural observation that only small changes took place in the geometry of the bridging His63 ligand relative to Zn in the Zn-containing subunit of the heterodimer upon reduction of Cu. The reduced WT SOD1 XANES spectrum was similar to that reported before (Hasnain et al., 1987), but the Zn XANES was markedly different from the D83S-C111S heterodimer indicating a shift in the electronic structure in the heterodimer consistent with long range changes in the WT monomer coordination environment around Zn due to the absence of Zn in the Zn-deficient subunit.

All Cu K-edge spectra for oxidized and reduced forms of D83S-C111S heterodimer and WT SOD1 (Fig. 6B) exhibit a peak at ∼8984 eV which is attributed to the 1s - 4p transition of Cu^I^(8980–8985 eV) (Kau et al., 1987; Streltsov and Varghese, 2008). The relatively high intensity of this pre-edge peak for reduced forms was consistent with a 3-coordinate Cu^I^ species (Blackburn et al., 1989; Kau et al., 1987). This is consistent with the bridging imidazole His 63 becoming protonated and the Cu moving to form a trigonal geometry with the remaining three His46, His48, His120 ligands. The reduced forms of both heterodimer and wild type SOD1 overlay quite well, suggesting that both may be reduced completely and the mutation in the zinc site only minimally affected the Cu^I^trigonal binding site. The lower heights of pre-edge peaks at ∼8984 eV for untreated form of heterodimer and wild type SOD1 suggest that mixed Cu oxidation states exist with high content of Cu^I^in the heterodimer.

The XANES data were collected and analyzed with the ATHENA package (Ravel and Newville, 2005) and Linear Combination Fitting (LCF) was used to estimate the reduction of copper in the heterodimer. Wild-type oxidized and reduced SOD1 were used as standards to approximate the complete oxidation and reduction of Cu. The XANES normalized µ(E) data converged to a Cu^I^:Cu^II^ratio of 0.74 / 0.26 (Fig. S2, Supplemental infromation). This corresponds to a Cu^I^/Cu^II^ ratio in the heterodimer of 2.8, which is close to the estimate of 2.6 based on structural refinement reported above and confirms the propensity of zinc-deficient SOD1 to be more easily reduced. The greater amount of the Cu^I^ reduced state in Zn-deficient D83S-C111S SOD1 heterodimer is consistent with the crystallographic fitting of increased Cu^I^/Cu^II^ in the x-ray structure and the increased chemical-reactivity measured in the peroxynitrite generation assay reported above.

## DISCUSSION

The tethering of SOD1 subunits allowed visualization of how the dimer interface of the Cu,Zn SOD1 subunit affects the zinc-deficient subunit. Remarkably, the structure of both subunits in the tethered heterodimeric SOD1 were essentially identical to the previously reported holo Cu,Zn SOD1 structure (Fig. 7A), with both the zinc-binding and the electrostatic loops in the zinc-deficient subunit assuming a native configuration. This structural order in the tethered zinc-deficient subunit is in sharp contrast to the disorder resulting from zinc missing in both SOD1 subunits. In the zinc-deficient homodimer, the zinc and electrostatic loops in both subunits are highly disordered with 11 and 8 residues respectively being unstructured in the structure (Fig. 7B). Because each zinc loop forms 30% of the dimer interface, the absence of zinc in both subunits understandably results in the dimer interface twisting by nine degrees (Fig. 7C). In addition, the disulfide bridge between C57-C146 is far more susceptible to reduction in the homodimeric zinc-deficient SOD structure (Roberts et al., 2007). As a consequence, homodimeric zinc-deficient SOD1 is more conformationally mobile than the heterodimer and therefore likely to further unfold, lose copper, and eventually undergo irreversible aggregation.

**Figure 7.**
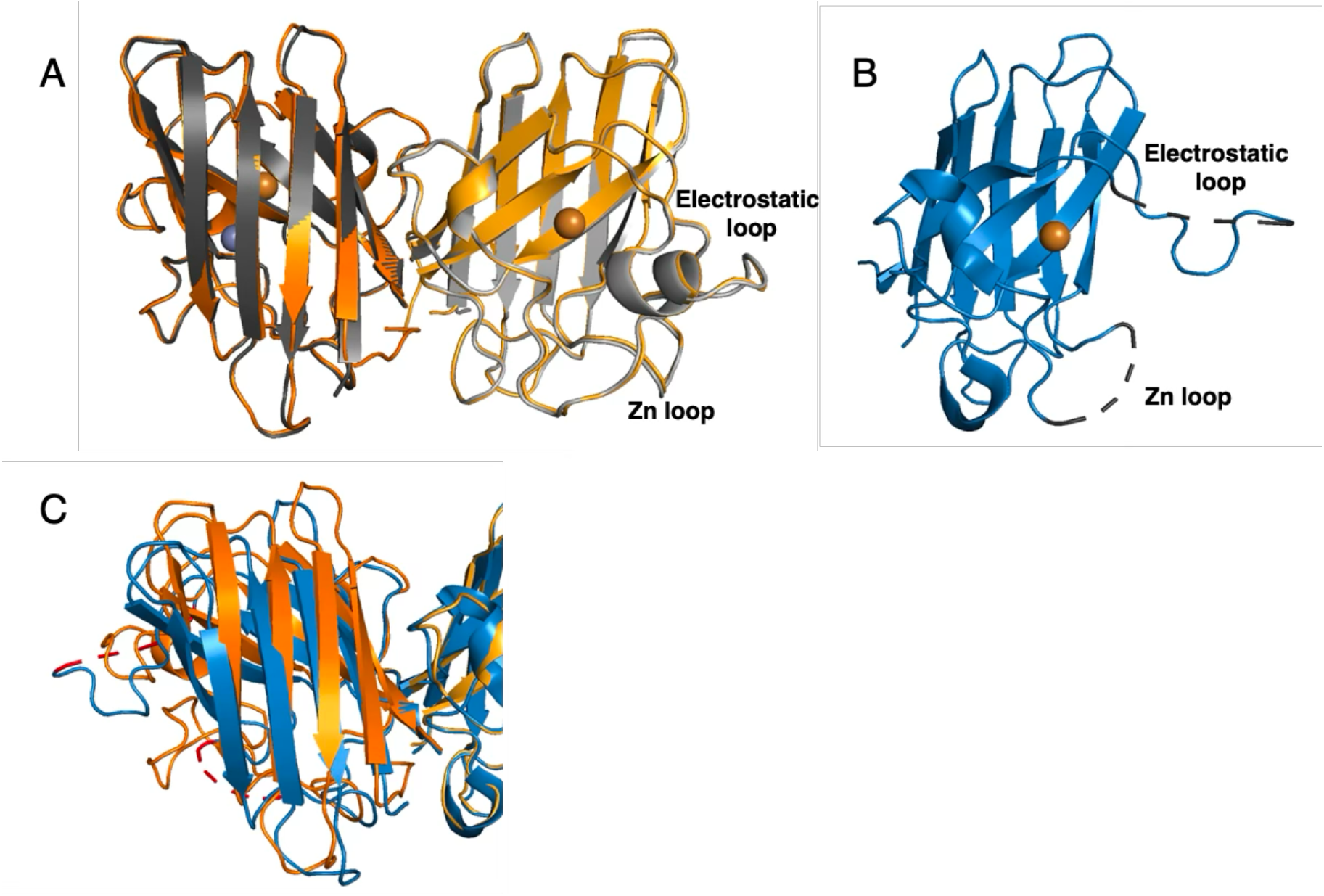
**A)**. Overlay of the tethered heterodimer SOD1 (orange) with WT SOD1 (grey). Only the metals bound to the tethered SOD1 are shown (Cu in orange and Zn in grey sphere) (Het-SOD1 PDB:6Dtk, WT SOD1 PDB:1PU0). **B.)** shows the electrostatic and zinc-binding loops disordered in zinc-deficient homodimer (blue, PDB:2R27) compared to the right zinc-deficient subunit in 7A the tethered heterodimer (16). **C.)** Overlay of tethered Het-SOD1 (orange) with the homodimeric zinc-deficient SOD1 illustrating the greater twist in the dimer interface in the zinc deficient-homodimer (blue).

The loss of zinc is generally recognized as an early and common intermediate in the unfolding of Cu,Zn SOD1 (Crow et al., 1997; Mulligan et al., 2008), leading to further misfolded species hypothesized to be toxic and prone to aggregation. However, the heterodimer, missing just one zinc, was itself not misfolded as shown in Fig 7A. Yet, the heterodimer was substantially more toxic to cultured primary motor neurons than the same amount of the zinc-deficient dimer (Fig. 3). The enhanced toxicity of the heterodimer reported here aligns with our previous results showing delivery of Cu, Zn SOD1 subunits mixed with zinc-deficient SOD1 were more damaging to motor neurons than the same amount of SOD1 that was entirely zinc-deficient (Garner et al., 2010). These results further align with multiple studies showing that transgenic mice overexpressing both wild-type SOD1 and familial ALS SOD1 suffer from an accelerated disease progression (Fukada et al., 2001; Jaarsma et al., 2000). Similarly, increased toxicity without aggregation has been reported for tethered SOD1 heterodimers in *C. elegans* (Witan et al., 2008). Together, these observations raise the intriguing possibility that the wild-type allele in human familial-SOD1 ALS patients can physically contribute to the dominant inheritance of SOD1 mutations through heterodimer formation.

Measuring zinc-deficient SOD1 in vivo has been a daunting task as the remaining copper is labile (Ellerby et al., 1996; Pantoliano et al., 1982) and may be lost during the ante-mortem interval and sample preparation. However, growing *in vivo* evidence now supports the metal loss from SOD1 with its malfunction in ALS. Zinc-deficient, copper-containing SOD1 has now been measured specifically in the ventral horn in autopsied spinal cord both sporadic and familial ALS patients (Trist et al., 2022). Notably, zinc-deficient SOD1 was not found in non-ALS control patient samples nor in non-disease affected CNS regions in the same patients. The zinc-deficient SOD1 was present mostly in a ratio of 1.67 copper atoms per zinc atom. This ratio is consistent with zinc-deficient SOD1 forming heterodimers with Cu, Zn SOD1.

Using native nano-DESI imaging mass spectrometry, Hale et al. (2025) have shown a striking localization of dimeric G93A-SOD1 missing one metal in the ventral horn of the spinal cord at an early symptomatic stage of motor neuron degeneration. The imaging of SOD1 with its bound metals provides strong evidence for a functional change in mutant SOD1 that is specifically associated with disease affected regions that is not present in transgenic WT SOD1-overexpressing mice. The resolution of the mass spectrometer could not distinguish whether G93A-SOD1 lost zinc or copper. However, the presence of dimeric three-metal G93A-SOD1 localized specifically in ventral spinal cord is suggestive that heterodimeric zinc-deficient SOD1 could be a significant toxic species and is present at disease onset in G93A SOD1 transgenic mice. The metal loss reported by Trist et al. (2022) and by Hale et al. (2025) provide important clues as to how the ubiquitous expression of mutant SOD1 throughout the whole body from birth can lead to the restricted degeneration of motor neurons in ALS.

The misfolding and aggregation hypotheses predict heterodimer formation would slow disease progression by decreasing aggregation. Conversely, the pro-oxidant SOD1 hypothesis (Garner et al., 2010) suggests heterodimers can help drive motor neuron degeneration by prolonging the lifetime of zinc-deficient SOD1. Our findings support the latter pro-oxidant SOD1 hypothesis. This challenges the idea that aggregation is the primary driver of toxicity in ALS (Benatar et al., 2025; Wang et al., 2008). Instead, the present results support the view that zinc-deficient SOD1 stabilized as a heterodimer increases toxicity and can contribute to the dominant inheritance of SOD1 mutations.

Thus, this work implicates WT Cu,Zn SOD1 being an active participant in the dominant gain-of-function responsible for motor neuron loss in ALS. The tethered D83S+WT SOD1 heterodimer recapitulated our previous findings that demonstrated Cu, Zn SOD1 greatly enhances the toxicity of zinc-deficient SOD1 (Drechsel et al., 2012; Garner et al., 2010). We further confirmed the mechanism for the enhanced motor neuron death by the tethered heterodimer was mediated a superoxide and nitric oxide-dependent oxidative mechanism as previously reported. We also used a boronate probe, which is a more selective detector for peroxynitrite (Zielonka et al., 2012), to further validate the generation of peroxynitrite during the reoxidation of SOD1. This oxidative death cascade catalyzed by zinc-deficient SOD1 has been elucidated in multiple prior studies over the past three decades (Beckman et al., 1993; Estevez et al., 2000; Estevez et al., 1998; Franco et al., 2013; Pehar et al., 2004; Raoul et al., 2002).

## Data availability statement

The data used and analyzed in this current study can be made available by the corresponding author upon reasonable request.

## ACKNOWLEDGMENTS

We acknowledge the use of the Australian Synchrotron Protein Crystallography (MX2) and X-ray Absorption Spectroscopy (XAS) beamlines, and the CSIRO Collaborative Crystallization Centre (C3), Victoria, Australia. Parts of this study were supported by a Bethlehem Griffiths foundation and Betty Laidlaw MND Research Grant. We would like to thank the Florey Institute of Neuroproteomics Facility. Funding provided by Emory start-up, National Health and Medical Research Council (NHMRC) project grant (APP1164692, APP1138673), National Institutes of Health (1R01AG085587, R01AG070937-03, U01AG061357-05). We also acknowledge funding from the Victorian Government’s Operational Infrastructure support program. The Emory Glycomics and Molecular interaction core (EGMIC, RRID:SCR0-023524) and the Ion, Omics and Neuroscience core (ION core).

