## Supplemental for "The structure, redox chemistry and motor neuron toxicity of heterodimeric zinc-deficient SOD1-Implications for the toxic gain of function observed in ALS"

**Supplemental Table 1. Source parameters for the 6545XT QTOF for native protein mass spectrometry.**

| **Agilent 6545XT AdvanceBio LC/Q-TOF** | |
|---|---|
| Source | Agilent Jet Stream |
| Dry Gas Temperature | 180 °C |
| Dry Gas Flow | 10 L/min |
| Nebulizer | 30 psig |
| Sheath Gas Temperature | 150 °C |
| Sheath Gas Flow | 10 L/min |
| VCap | 5000 V |
| Nozzle Voltage | 2000 V |
| Fragmentor | 300 V |
| Skimmer | 220 V |
| Quad AMU | *m/z* 500 |
| Mass Range | *m/z* 300–7000 |
| Acquisition Rate | 1.0 spectrum/s |
| Acquisition Mode | Positive, extended (*m/z* 10,000) mass range |

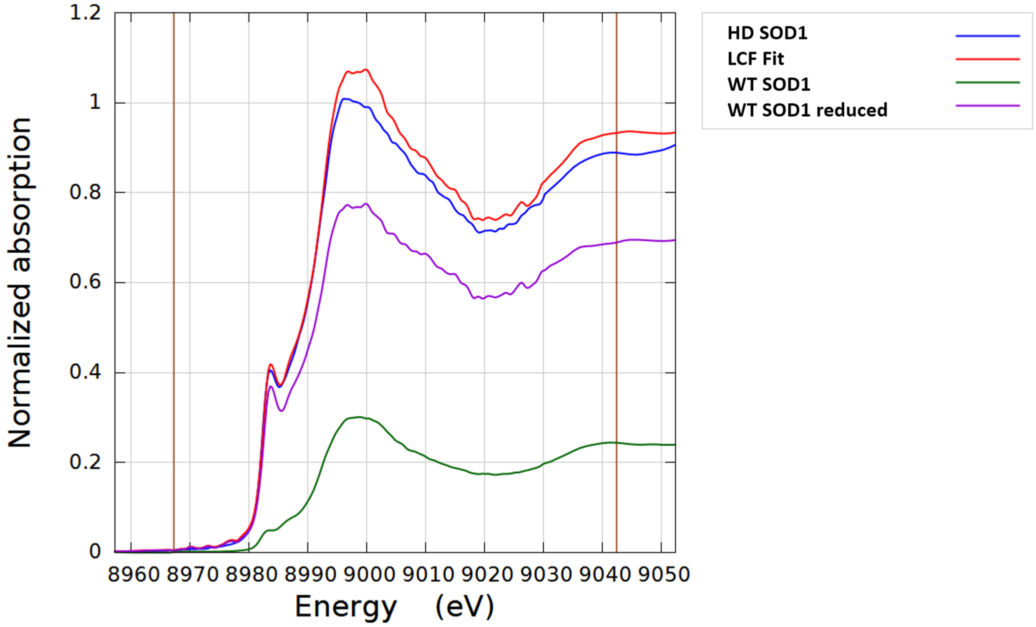


**Figure S1.** Linear combination fitting (LCF) to the D83S-C111S SOD1 heterodimer XANES data using WT and reduced SOD1 data as the fitting standards. LCF is based on XANES normalized μ(E) data region ($E_0$ -15, + 60 (eV); $E_0$ – the photoelectron energy threshold) marked by vertical lines.
